# Oral administration of bacterial Beta cell expansion factor A (BefA) alleviates diabetes in mice with Type 1 and Type 2 diabetes

**DOI:** 10.1101/2021.10.22.465460

**Authors:** Huan Wang, Jing Wei, Hong Hu, Fuyin Le, Heng Wu, Hong Wei, Jie Luo, Tingtao Chen

## Abstract

Diabetes mellitus (DM) is a group of metabolic diseases, which is of urgent need to develop new therapeutic DM oral drugs with less side effects and sound therapeutic efficacy. In this study, a Beta cell expansion factor A (BefA) production strain of *Escherichia Coli* BL21-pet 28C-BefA was constructed, and the anti-diabetes effect of BefA was evaluated using type 1 diabetes mellitus (T1DM) and type 2 diabetes mellitus (T2DM) mice models. The T1DM mice results indicated that BefA significantly reduced the blood glucose level, exerted protective function of islet β cell morphology, down-regulated the TLR-4, p-NFκB/NFκB, Bax/Bcl-2 expressions and the secretion level of IL-1β, TNF-α, increased the expression of PDX-1 protein and insulin secretion in a concentration-dependent manner, and restored the disturbed microbial diversity to normal level. Similar with the T1DM mice, BefA obviously increased islet β cells, reduced inflammatory reaction and apoptosis in T2DM mice, and also improved liver lipid metabolism by down-regulating the expression of CEBP-α, ACC, Fasn and inhibiting the synthesis of triglyceride and induce Cpt-1, Hmgcs2, Pparα in a concentration-dependent manner. In the present study, we verified therapeutic effect and potential mechanisms of BefA in mammal for the first time, providing basic data for its clinical application.

## Introduction

Diabetes mellitus (DM) is a group of complex metabolic disorder characterized by abnormally elevated blood glucose concentrations secondary to either insufficient insulin secretion, insulin resistance, or both (Association, 2012; Weir and Bonner-Weir, 2013). Studies show that acute complications, e.g. hyperosmolar coma and diabetic ketoacidosis, can be caused by elevated blood glucose, which eventually cause damage to liver, heart, cerebrovascular and other organs and even lead to death (Sattar, 2013; Kumar et al., 2017). The global mortality rate of DM is as high as 10.7%, and it is estimated that there will be 693 million people with DM worldwide by 2045 (Cho et al., 2018). The increasing prevalence of DM and its high medical expenses make it an urgent public health problem all over the world (Zimmet et al., 2014; Iwasaki et al., 2016).

Insulin-dependent type 1 diabetes mellitus (T1DM) and insulin-independent type 2 diabetes mellitus (T2DM) are the main types of diabetes, among which T1DM results from the specific deficiency of insulin-producing pancreatic β cell from autoimmune destruction (Jahromi and Eisenbarth, 2007), and T2DM is an age-related disease characterized by the dis-function of glucose metabolism representing insulin-resistant states, accompanied by a destruction of β cell function (Weir and Bonner-Weir, 2013). For the treatment of T1DM, insulin injection therapy was applied after its discovery in 1922, and it can only alleviate but fail to eliminate T1DM, and also may cause long-term physical suffering through subcutaneous injection (Amiel and Rela, 2005). New methods such as immunotherapy, gene therapy, and organ transplantation have been developed rapidly, but they are still in the research stage due to problems such as therapeutic side-effects, safety issues or insufficient donors (Song et al., 2015; Song and Roy, 2016; Im and Bhang, 2018). For the treatment of T2DM, various drugs were developed but people found their defects during clinical practice. For example, biguanide drugs like metformin can lead to macrocytic anemia and increase the burden of liver and kidney (Inagaki et al., 2016); sulfonylurea drugs can enhance insulin sensitivity as well as cause digestive system disease and liver function impairment(Hsu et al., 2011); new drugs such as GLP-1 receptor agonists has remarkable curative effect but the expensive cost and the need for injection limit its clinical use(Wit et al., 2016). Based on the disadvantages of the above treatment strategies, it is of great importance to develop new therapeutic T1DM and T2DM drugs with less side effects and better therapeutic efficacy. The impaired pancreatic β cells function and insulin secretion have been demonstrated in both T1DM and T2DM, while treatments targeting pancreatic β cell proliferation are currently lacking (Elizabeth et al., 2017; Mirmira et al. 2016).

Many studies proved that intestinal microbes and their metabolite exert important effects on obesity and blood glucose metabolism, but the direct evidence is not clear. In 2016, a research published on *eLife* reported an intestinal microbiota-derived protein named BefA (Beta cell expansion factor A) that could induce pancreatic β cell proliferation in early development of zebrafish. More meaningfully, the research team discovered that the BefA protein homologues in human intestinal microbial metabolites sharing the same proliferative effect. Besides, since BefA protein is derived from intestinal microbes so it possesses high tolerance to intestinal environment than other drugs, and can be administered orally to avoid the physiological pain caused to patients by repeated injections, indicating that BefA protein may be a new strategy for DM treatment (Hill et al., 2016).

Pancreatic islet β cells of T1DM mice will be irreversibly destroyed by abnormal autoimmune attack under normal pathological conditions (Tai et al., 2016), some key proteins are associated with apoptosis, such as PDX-1 (Pancreas/duodenum homeobox protein 1, the marker of islet β cell differentiation, maturation and proliferation) (Oliver-Krasinski et al., 2009; Wang et al., 2020). Similarly, Mafa (V-maf musculoaponeurotic fibrosarcoma oncogene homolog A) and Neurogenin-3 are critical transcription regulators in inducing β cell development and regeneration (Zhu et al., 2017). Furthermore, chronic systemic inflammation plays a promoting role in the occurrence and development of T2DM (Badawi, 2011), the key inflammatory proteins includes TLR-4, p-NFκB/NFκB, IL-1β and TNF-α.

Based on the irreversible damage of pancreatic β cells in T1DM and T2DM,this work shed light on the role of the increasing effect of BefA protein by adjusting blood glucose level, body weight and gene expression changes associated with β cell proliferation and necrosis. In addition, as the disordered intestinal microbiota can lead autoimmune damage on islet β cells and induce T1DM (Wen et al., 2008), and T2DM is prone to occur in obese individuals and the high fat diet could induce fatty liver and liver inflammation (Jiao et al., 2010), so we also evaluated the effect of orally administration of BefA on intestinal microbiota and liver function. In the present study, we construed the BefA yield strain, isolated and purified the BefA protein, and evaluated its therapeutic effect and potential mechanisms in T1DM and T2DM mice for the first time, providing basic data for its clinical application.

## Results

### Increasing effect of purified BefA on the number of islet β cell in newborn mice

To obtain the purified BefA, the codon optimized BefA gene was inserted into the prokaryotic expression vector pet 28C, to make the BefA production strain of BL21-pet 28C-BefA. As shown in Fig. 1a, massive BefA produced by BL21-pet 28C-BefA strain was presented as soluble protein, and the purity of soluble BefA protein could reach above 98%, and their accurate (25-28 kD in size) was further confirmed using Western-blotting results (Fig. 1B).

**Figure 1.**
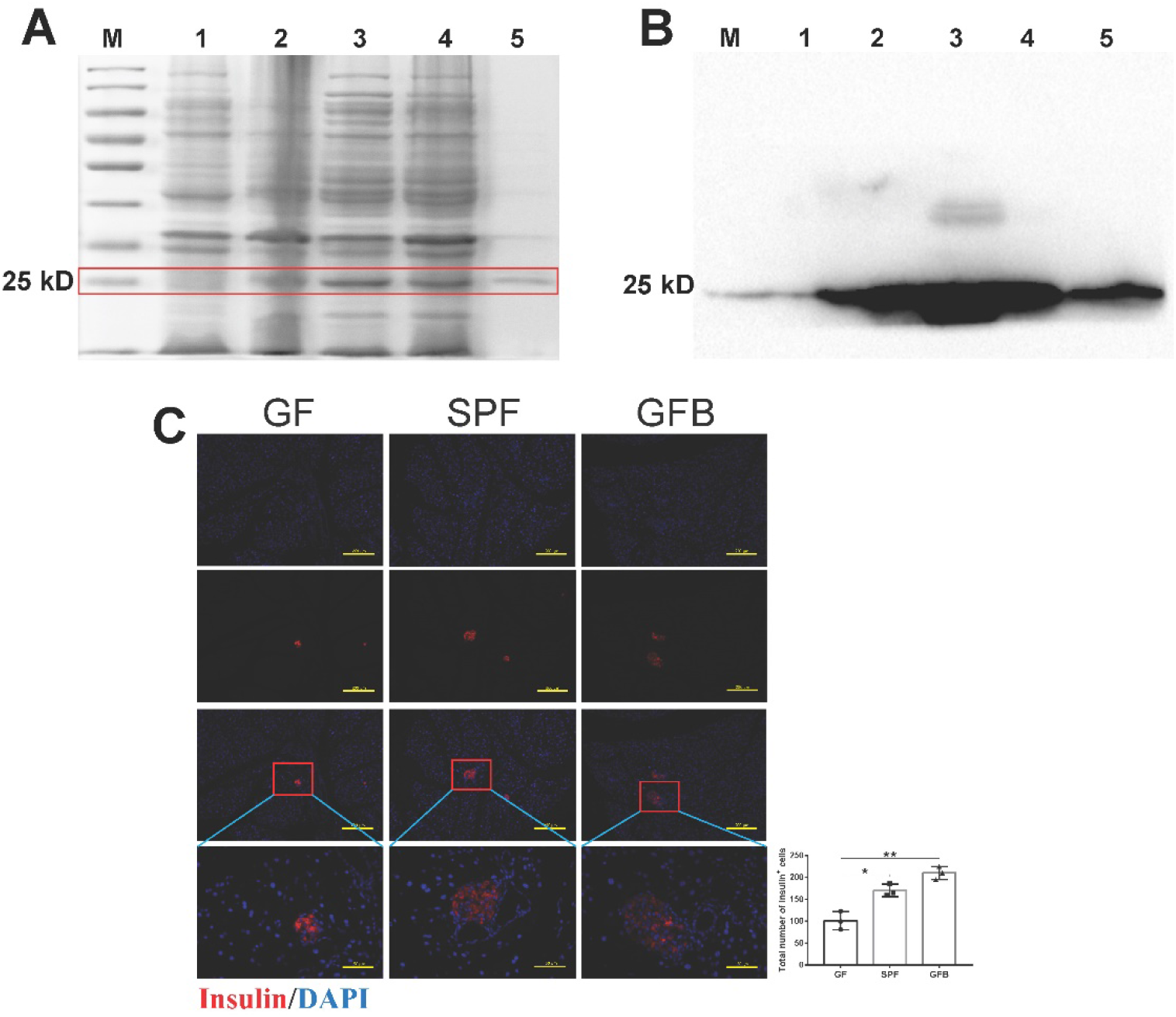
Increasing effect of purified BefA on the number of islet β cell in newborn mice. (A) Protein purity detection using SDS-PAGE. (B) Verifying the band position of the BefA protein using Western-blotting. (C) BefA increased the number of islet β cells in GF mouse. The red area marks secreted insulin, which is used to label islet β cell. M: protein marker. 1: Bacteria after IPTG induction. 2: Bacterial lysate supernatant. 3: Bacterial lysate precipitation. 4: Supernatant purified protein. GF: germ free mouse (n = 3). SPF: specific pathogen free mouse (n = 3). GFB: germ free mouse + BefA protein treatment (n = 3, 1 ng BefA/g body weight of BefA).

As no work is done to verify the increasing effect of orally administered BefA on the number of mammalian islet β cells, therefore newborn germ-free (GF) mice and specific pathogen free (SPF) mice were used in the present study to confirm the anti-diabetes effect of BefA for the first time, and our results indicated that BefA could markedly increase the β cell number compare with SPF mice and GF mice (Fig. 1C), consistent with previous work in zebrafish (Hill et al., 2016).

### BefA protein significantly reduced pancreatic inflammation in T1DM mice

To evaluate the therapeutic effect of BefA in the mammalian model for diabetes, we established mice model of T1DM divided into 4 groups including C group (normal control group), M group (T1DM model group), MB10 group and MB50 group (Fig. 2A). As shown in Fig. 2B, 10 μg BefA and 50 μg BefA could significantly reduce the blood glucose levels in a concentration-dependent manner (p < 0.05, Fig. 2B), and had little effect on body weight (Fig. 2C).

**Figure 2.**
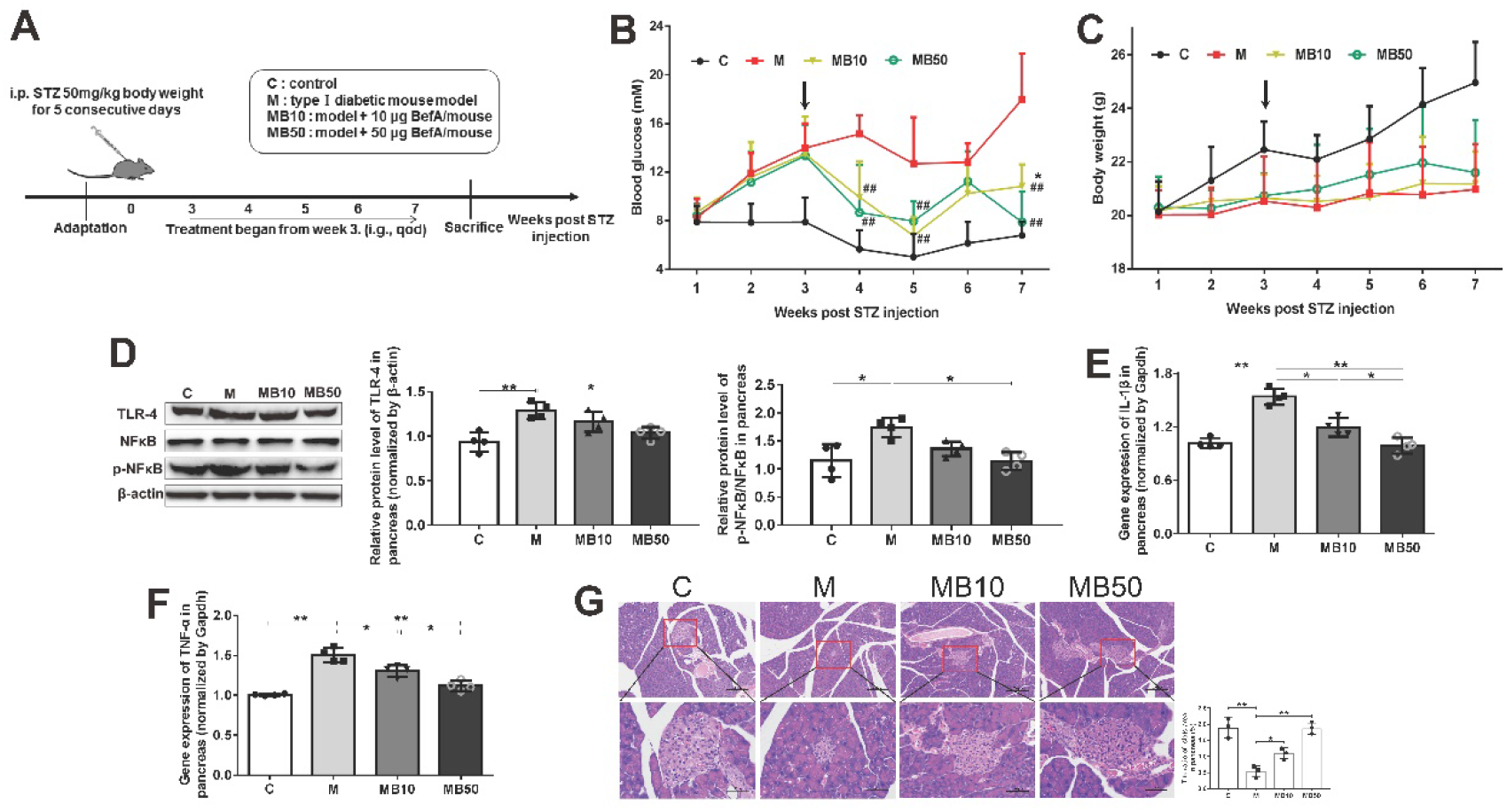
BefA protein significantly reduced pancreatic inflammation in T1DM mice. (A) Scheme of animal experiment. (B) Blood glucose level tested from mice tail vein. (C) Body weight. Western-blotting results of the expression level of pro-inflammatory proteins including TLR-4 and the ratio of p-NFκB/NFκB in T1DM mice pancreas (D), gene expression levels of IL-1β (E) and TNF-α (F) in pancreas measured by q-PCR. (G) Results of HE staining (100 × and 400 ×). The black arrows in (A) and (B) indicated that the BefA administration time started from the week 3. C group: wild-type C57BL/6 mice (n = 15); M group: T1DM model group treated with 0.9% physiological saline containing 0.01% gelatin administered intragastrically every other day for 14 times (n = 15); MB10 group: T1DM model group treated with 0.9% physiological saline containing 0.01% gelatin and 10 μg BefA administered intragastrically every other day for 14 times (n = 15); MB50 group: T1DM model group treated with 0.9% physiological saline containing 0.01% gelatin and 50 μg BefA administered intragastrically every other day for 14 times (n = 15). Data are presented as mean ± SD. From (B-C), * means significant difference compared with C group, p < 0.05; ## means significant difference compared with M group, p < 0.01. From (d-f), * p < 0.05, ** p < 0.01.

As the damage of islet β cell has a strong connection with pancreatic inflammation, we further studied the effect of BefA on inflammatory signaling pathway, and our results indicated that 50 μg BefA significantly down-regulated the TLR-4 expression and p-NFκB expression (p < 0.05, Fig. 2D), and both 10 μg BefA and 50 μg BefA had effectively reduced the production of IL-1β and TNF-α at transcription level (p < 0.05, Fig. 2E and F). The H&E staining of mice pancreas further confirmed that BefA protein could obviously increase the islet β cells in a concentration-dependent manner (Fig. 2G).

### BefA protein could reduce pancreas injury and restore intestinal microbiota to normal level in T1DM mice

To evaluate the effect of BefA on the pancreas of T1DM mice, some key proteins associated with apoptosis were tested. As shown in Fig. 3A, injection of STZ obviously induced the apoptosis in pancreas, and the use of 10 μg BefA and 50 μg BefA significantly reduced the cell necrosis in pancreas by 48% and 66% compared with M group, respectively (p < 0.01), and BefA protein also significantly enhanced the expression of PDX-1 compared with M group (p < 0.01, Fig. 3B and C) and increased the insulin secretion level in a concentration-dependent manner (p < 0.05, Fig. 3D).

**Figure 3.**
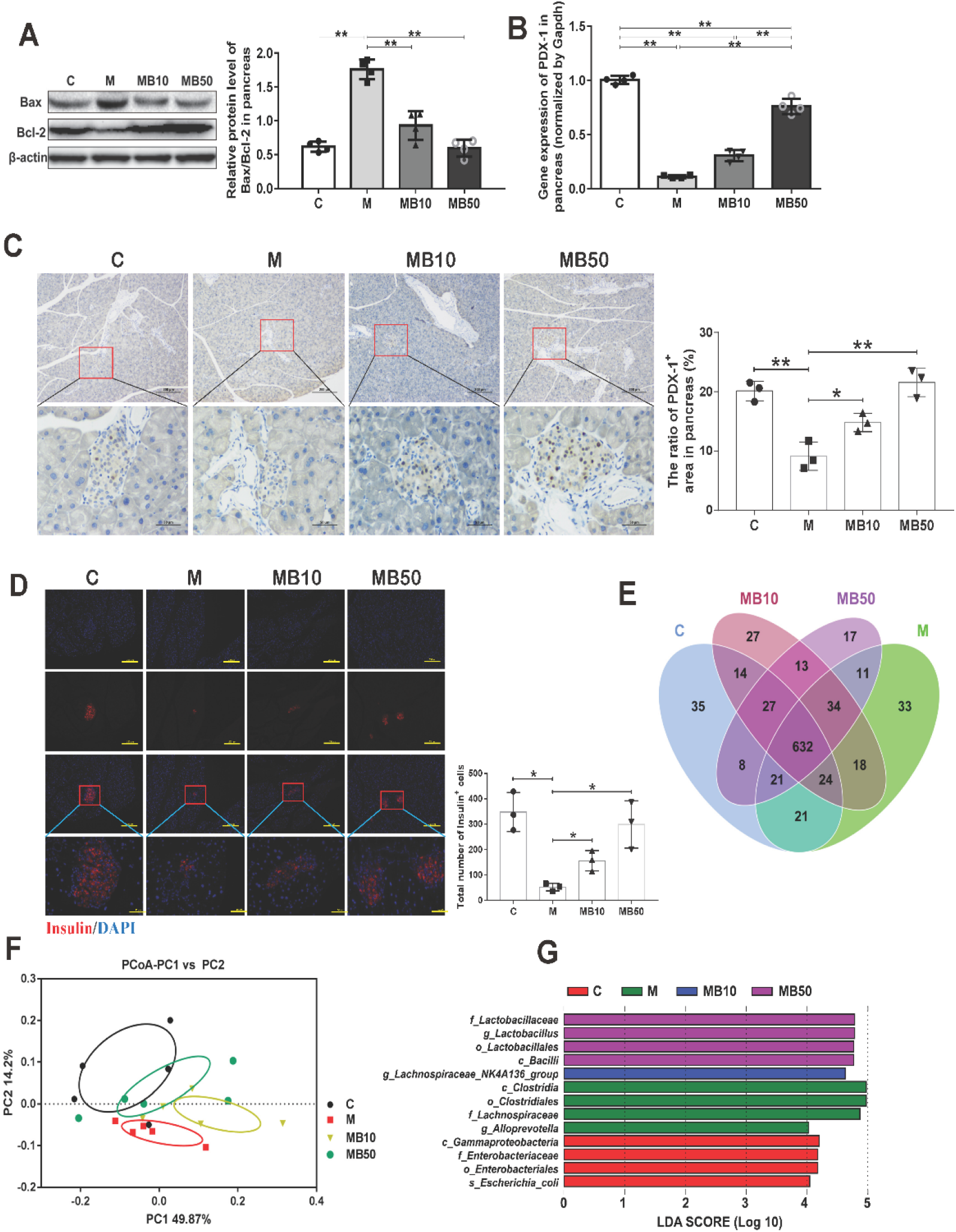
BefA protein could reduce pancreas injury and restore intestinal microbiota to normal level in T1DM mice. (A) Western-blotting results of the expression ratio of Bax/Bcl-2. (B) q-PCR results of the transcription level of PDX-1 protein. (C-D) The immunohistochemistry analysis of PDX-1 protein (100 × and 400 ×) and the immunofluorescence analysis of insulin (100 × and 400 ×) in pancreas. The red area marks secreted insulin, which is used to label islet β cell. High-throughput sequencing results including Venn diagram (E), PcoA analysis (F) and LEfSe analysis (G) of T1DM mouse feces among different groups. C group: wild-type C57BL/6 mice (n = 15); M group: T1DM model group treated with 0.9% physiological saline containing 0.01% gelatin administered intragastrically every other day for 14 times (n = 15); MB10 group: T1DM model group treated with 0.9% physiological saline containing 0.01% gelatin and 10 μg BefA administered intragastrically every other day for 14 times (n = 15); MB50 group: T1DM model group treated with 0.9% physiological saline containing 0.01% gelatin and 50 μg BefA administered intragastrically every other day for 14 times (n = 15). Data are presented as mean ± SD. * p < 0.05, ** p < 0.01.

In the analysis of the effect of BefA on intestinal microbiota by using high-throughput sequencing, the Venn diagram indicated that 632 OTUs were determine to be common OTUs among all groups, accounting for 80.82% (C group, 632/782), 79.60% (M group, 632/794), 80.10% (MB10 group, 632/789) and 82.83% (MB50 group, 632/763), respectively (Fig. 3E). The PCoA result revealed a closer distance between samples in C group and MB50 group (Fig. 3F), indicated that the BefA have a positive effect to restore intestinal microbial composition to normal level, charactering by the high abundance of probiotic *Lactobacillus* at family (f), genus (g) and order (o) levels than other three groups (Fig. 3G).

### BefA protein showed sound protective effect on pancreas in T2DM mice

To explore the anti-T2DM effect of BefA in mammals, a T2DM mice model was established and divided into 5 groups including C group (normal control group), M group (T2DM model group), MB5 group, MB10 group and MB40 group (Fig. 4A), and the effect of BefA on the body weight, blood glucose and glucose tolerance of mice were tested. The results indicated that 40 μg BefA could significantly reduce the weight gain symptoms of T2DM mice (p < 0.05, Fig. 4B), and both 20 μg BefA and 40 μg BefA could reduce the blood glucose level, of which a 41% reduction in MB40 group was obtained at week 10 compared with that in M group (p < 0.01, Fig. 4C). On 10^th^ week, oral glucose tolerance test was performed and 40 μg BefA had a significant improvement of glucose tolerance compared with M group since 60 min after glucose injection (p < 0.05, Fig. 4D).

**Figure 4.**
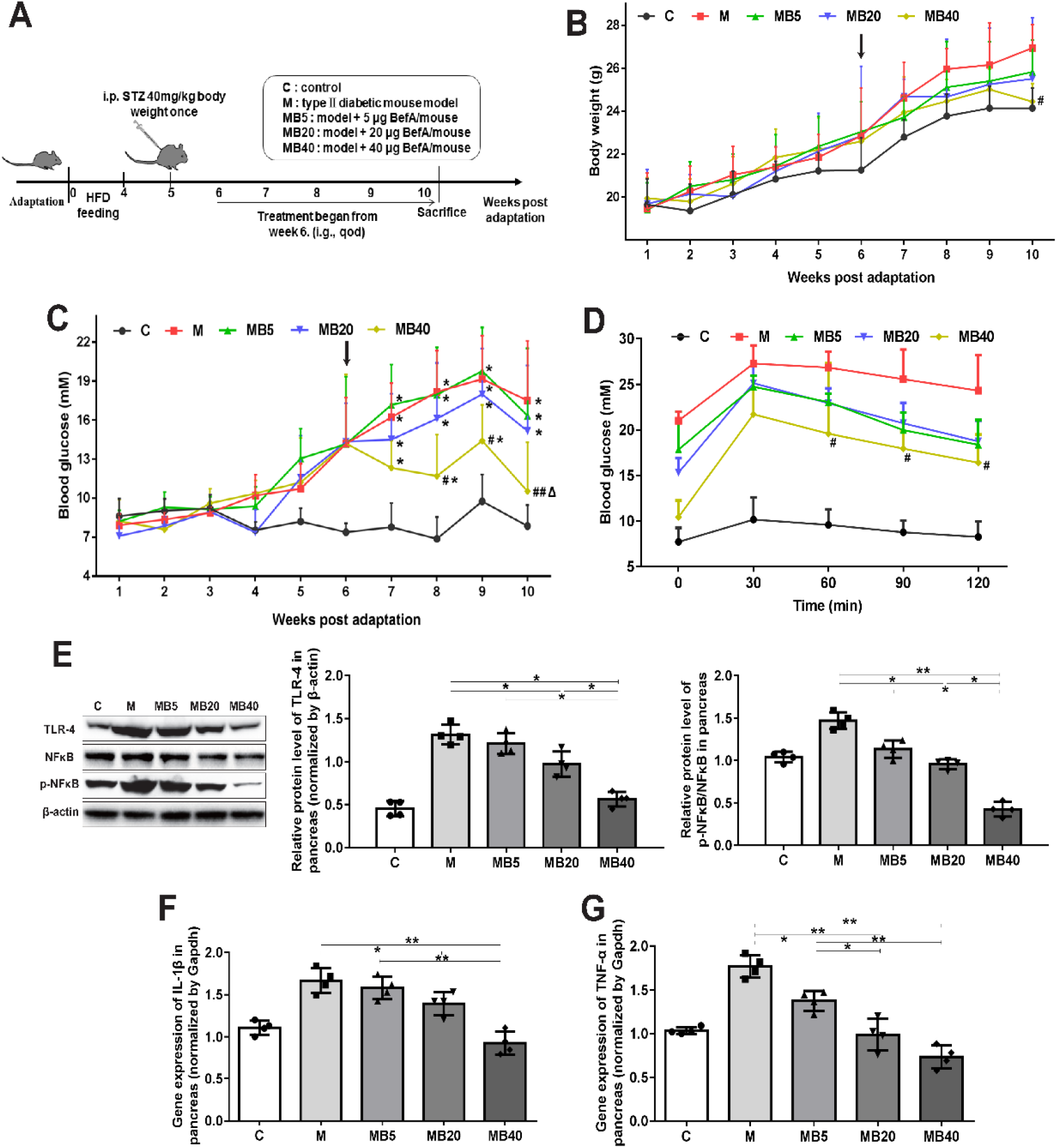
BefA protein showed sound protective effect on pancreas in T2DM mice. (A) Scheme of animal experiment (n = 15). (B) Body weight. (C) The blood glucose levels from mouse tail vein tested every week. (D) OGTT test performed on 10th week. (E) Western-blotting results of the expression level of pro-inflammatory proteins including TLR-4 and the ratio of p-NFκB/NFκB in T2DM mouse pancreas. Gene expression levels of IL-1β (F) and TNF-α (G) in pancreas measured by q-PCR. The black arrows in (B) and (C) indicated that the BefA administration time started from the week 6. C group: mice fed with laboratory chow diet as normal control group; M group: mice treated with 0.9% physiological saline containing 0.01% gelatin administered intragastrically every other day for 14 times (n = 15); MB5 group: mice treated with 0.9% physiological saline containing 0.01% gelatin and 5 μg BefA (n = 15); MB20 group: mice treated with 0.9% physiological saline containing 0.01% gelatin and 20 μg BefA (n = 15); MB40 group: mice treated with 0.9% physiological saline containing 0.01% gelatin and 40 μg BefA (n = 15). Data are presented as mean ± SD. * p < 0.05, ** p < 0.01.

Due to the important role of chronic systemic inflammation in the occurrence and development of T2DM, several key inflammatory proteins were detected. As shown in Fig. 4E-G, key inflammatory factors including TLR-4, p-NFκB/NFκB, IL-1β and TNF-α in pancreas were significantly down-regulated by BefA protein in a concentration-dependent manner. Compared with M group, 40 μg BefA significantly reduced the Bax/Bcl-2 ratio from 1.55 to 0.43 (p < 0.01, Fig. 5A) as well as increased the PDX-1 transcription level from 0.43 to 1.01 (p < 0.01, Fig. 5B), which was consistent with PDX-1 expression level in Fig. 5C. Mafa and Neurogenin-3 also showed a concentration-dependently upward expression trends in Fig. 5B. In addition, immunofluorescence test targeting insulin showed that BefA protein had a sound effect to promote insulin secretion level in a concentration-dependent manner (Fig. 5D).

**Figure 5.**
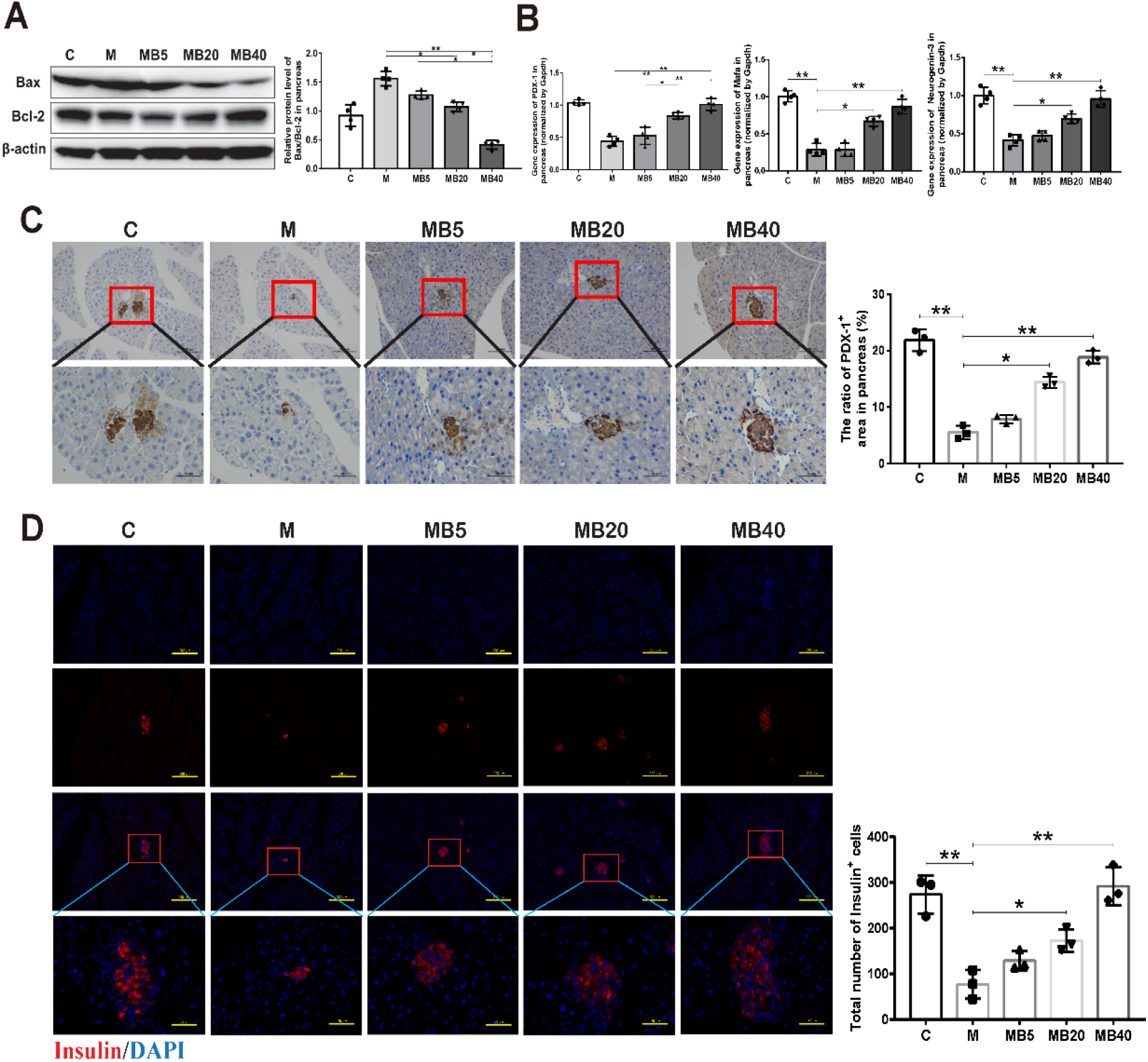
BefA protein showed sound effect on increasing the number of islet β cell and insulin secretion in T2DM mice. (A) Western-blotting results of the expression ratio of Bax/Bcl-2 in pancreas. The gene transcription and protein expression levels of PDX-1 performed by q-PCR (B) and immunohistochemistry analysis (C, 200 × and 400 ×). (D) Results of pancreatic immunofluorescence staining targeting insulin among different groups (100 × and 400 ×). C group: mice fed with laboratory chow diet as normal control group; M group: mice treated with 0.9% physiological saline containing 0.01% gelatin administered intragastrically every other day for 14 times (n = 15); MB5 group: mice treated with 0.9% physiological saline containing 0.01% gelatin and 5 μg BefA (n = 15); MB20 group: mice treated with 0.9% physiological saline containing 0.01% gelatin and 20 μg BefA (n = 15); MB40 group: mice treated with 0.9% physiological saline containing 0.01% gelatin and 40 μg BefA (n = 15). Data are presented as mean ± SD. *p < 0.05, ** p < 0.01.

### BefA protein showed sound effect on regulating liver lipid metabolism in T2DM mice

Because of the high occurrence of fatty liver and liver inflammation in T2DM patients, we evaluated the effect of BefA on liver function. The results indicated that BefA significantly down-regulated the expression level of CEBP-α (CCAAT/enhancer binding protein-α) a protein promoting adipocyte differentiation and accelerate fat accumulation, from 0.64 in M group to 0.25 in MB 40 group (p < 0.01, Fig. 6A), and significantly reduced the CEBP-α expression at the transcription level in a concentration-dependent manner (Fig. 6B). Moreover, the liver oil red O staining results indicated that a better fat-reducing effect was observed in higher concentration of BefA group (p < 0.01, Fig. 6C). In Fig. 6D, we found that BefA increased the mRNA expression of genes involved in the regulation of β-oxidation of fatty acids, including Carnitine palmitoyltransferase 1 (Cpt-1), 3-hydroxy-3-methylglutaryl-CoA synthase 2 (Hmgcs2) and Peroxisome proliferator-activated receptor α (Pparα), and inhibited the mRNA expression of lipogenesis-associated genes including Acetyl-CoA Carboxylase (ACC) and Fatty acid synthase (Fasn) in a concentration-dependent manner. In addition, BefA protein also significantly down-regulated levels of key inflammation proteins (TLR-4, p-NFκB/NFκB, IL-1β and TNF-α) in Fig. 6E-G (p < 0.05).

**Figure 6.**
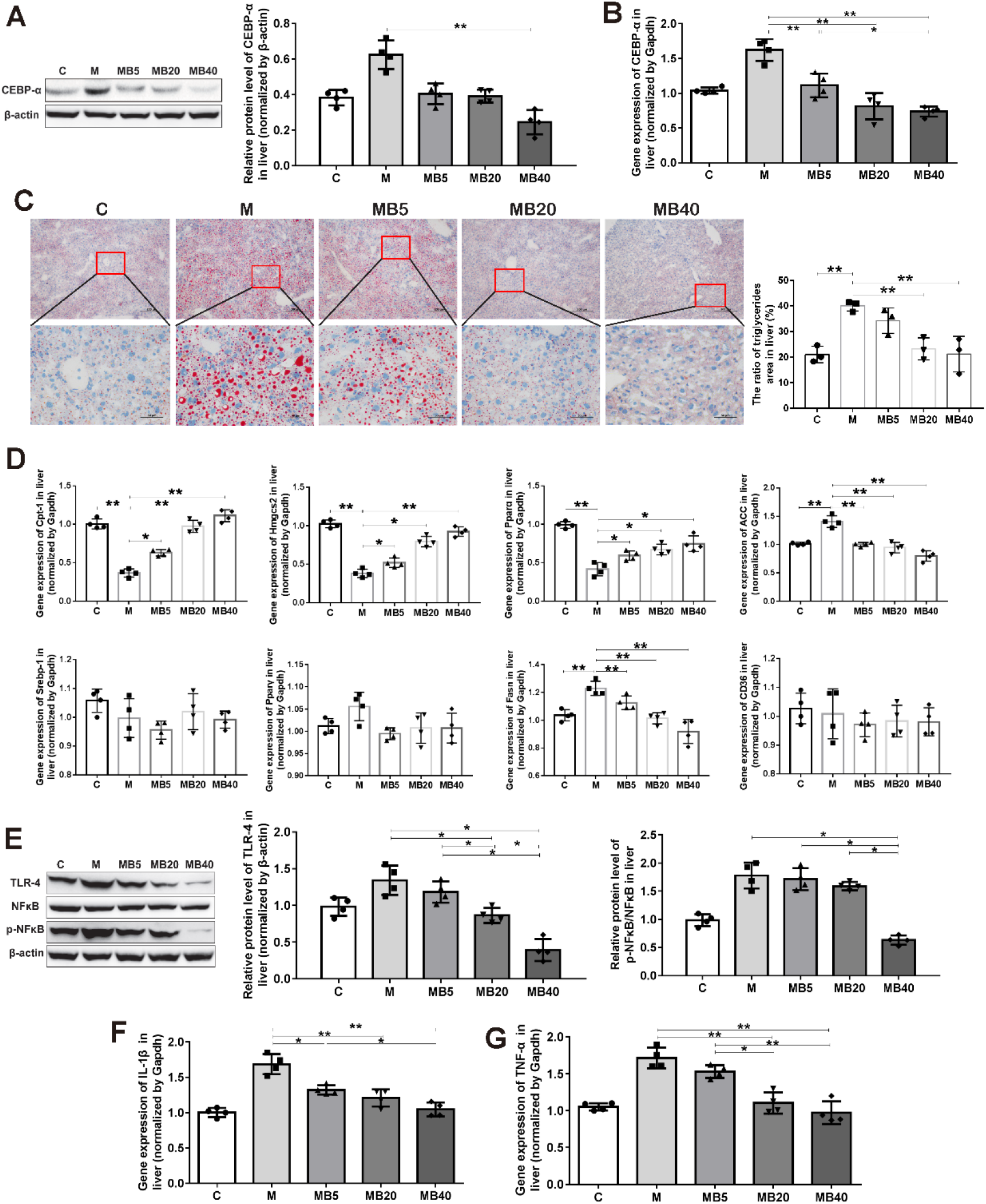
BefA protein showed sound effect on regulating liver lipid metabolism in T2DM mice. (A-B) The change of the expression and transcription level of CEBP-α gene measured by western-blotting and q-PCR. (C) Liver oil red O staining (100 × and 400 ×). (D) q-PCR results of lipid metabolism related factors including Cpt-1, Hmgcs2, Pparα, ACC, Srebp-1, Pparγ, Fasn and CD36 in liver (n = 4). (E) Western-blotting results of the expression level of pro-inflammatory proteins including TLR-4 and the ratio of p-NFκB/NFκB in T2DM mouse livers. Gene expression levels of IL-1β (F) and TNF-α (G) in livers measured by q-PCR. C group: mice fed with laboratory chow diet as normal control group; M group: mice treated with 0.9% physiological saline containing 0.01% gelatin administered intragastrically every other day for 14 times (n = 15); MB5 group: mice treated with 0.9% physiological saline containing 0.01% gelatin and 5 μg BefA (n = 15); MB20 group: mice treated with 0.9% physiological saline containing 0.01% gelatin and 20 μg BefA (n = 15); MB40 group: mice treated with 0.9% physiological saline containing 0.01% gelatin and 40 μg BefA (n = 15). Data are presented as mean ± SD. *p < 0.05, ** p < 0.01.

## Discussion

DM is a chronic systemic metabolic disease caused by long-term genetic and environmental factors (Saeedi et al., 2019). Although pathological study of DM concerning genetics, immunology and endocrinology has been developed in recent years, preventive and therapeutic methods still remain limited with drug resistance, side effects, suffering administration method as well as high economic burden cause negative influence in treatment effects and patients’ life quality (Ewald and Philip, 2013; Song and Roy, 2016; Kheirandish et al., 2017). Therefore, it is of great importance to develop new DM drugs with minor side effects and low economic cost.

Although the causes and pathological features of T1DM and T2DM are different, they also have common features of the irreversible damage on islet β cells (Mirmira et al., 2016; Elizabeth et al., 2017). Therefore, the drug can delay the occurrence of T1DM and T2DM directly from the origin if it can promote islet β cells proliferation. Previous study indicated that BefA could promote the proliferation of juvenile islet β cells in zebrafish, and BefA homologs was confirmed to be existed in human intestine that exert the same function (Hill et al., 2016), which showed a potential anti-diabetes of BefA in mammalian DM.

Firstly, we generated the BL21-pet 28C-BefA strain to produce BefA, and verified the increasing effect of BefA on the amount of mammalian islet β cell for the first time using GF mice (Fig. 1). Then, the T1DM mice model induced by STZ was established, STZ could increase reactive oxygen species (ROS) to accelerate islet β cell DNA damage and directly destroyed islet β cells, sharing typical human T1DM symptoms (Radenković et al., 2016). As β-cell damage mainly occurs in the islet inflammation area and TLR-4/NFκB is a pro-diabetic inflammation pathway, whose high expression will trigger the release of inflammatory factors IL-1β and TNF-α, and pro-inflammatory factors will inhibit the function of β cells and enhance the cytotoxicity, which eventually lead to a large number of islet β cells irreversible damage (Yang et al., 2018). Consistent with results in zebrafish, BefA significantly reduced the blood glucose level, exerted protective function of islet β cell morphology and cell density, down-regulated the expression level of TLR-4, p-NFκB/NFκB and the secretion level of IL-1β, TNF-α in a concentration-dependent manner (Fig. 2). T1DM will undergo abnormal islet β cell apoptosis, therefore we tested the expressions of Bax/Bcl-2 and PDX-1 in pancreatic tissue. Bax and Bcl-2 are classified as members of Bcl-2 family, among which Bax is up-regulated in apoptosis, and Bcl-2 is an important anti-apoptotic protein, and the ratio Bax/Bcl-2 is often used to evaluate their combined effect (Tomita, 2017). PDX-1, also called insulin promotor 1, is considered as an irreplaceable transcriptional factor in the differentiation and proliferation of islet β cells (Holland et al., 2005). The results indicated that BefA could recover the number of islet β cell and islet function of T1DM mice via reducing the Bax/Bcl-2 ratio, increasing the expression of PDX-1 protein and promoting insulin secretion (Fig. 3).

More and more studies indicated that intestinal microbiota was closely related to the development of diabetes (Hu et al., 2015; Paschou et al., 2018), so we conducted high-throughput sequencing analysis of intestinal microbiota in T1DM mice. The results showed that BefA could restore the disturbed microbial composition in M group to normal level, and markedly increased the abundance of probiotic lactic acid bacteria (LAB) in MB50 group (Fig. 3). LAB, such as *Lactobacillus, Bifdobacterium* can inhibit pathogenic bacteria, protect the barrier function of intestinal wall to protect the health of the body (Calix-Lara et al., 2014; Li et al., 2016). Moreover, LAB can decrease blood glucose level by directly decomposing glucose as well as reducing the expression of glucose transporter (GLUT) protein and inhibiting the glucose absorption in intestinal wall (Wang, 2020).

In T2DM mice, BefA also obviously reduced the blood glucose level as well as relieve weight gain symptoms in a concentration-dependent manner (Fig. 4). Notably, the insulin resistance, which is an important pathological indicator for evaluating glucose metabolism ability and insulin sensitivity in T2DM individuals (Saini, 2010), were significantly improved in MB40 group, indicating that BefA protein played a positive role in regulating blood glucose metabolism. Similar with the results in T1DM mice model, oral administration of BefA could reduce pancreatic inflammation by down-regulating the expression of TLR-4, p-NFκB/NFκB, IL-1β, TNF-α as well as promote islet β cell proliferation by lowering cell apoptosis (Bax/Bcl-2) and increase the expression of PDX-1 (Fig. 4, Fig. 5). In addition, the results suggest that BefA can promote β-oxidation of fatty acids by up-regulating of Cpt-1, Hmgcs2, Pparα and inhibit lipogenesis by down-regulating the expression of CEBP-α, ACC, Fasn, as well as inhibiting the synthesis of triglyceride, leading to improvement of fatty liver and metabolic dysfunction in a concentration-dependent manner (Fig. 6). Patients with T2DM can induce various complications including fatty liver and liver inflammation, dysfunction of hepatic lipid metabolism could be improved by activation of Cpt-1, Hmgcs2 and Pparα to enhance hepatic oxidation and metabolism of adipose tissue (Vilà-Brau et al., 2011; Montagner et al., 2016; Lundsgaard et al., 2018). CEBP-α, ACC, Fasn play an important role in initiating liver fat formation and lead to fatty liver (Horton et al., 1998; Jensen-Urstad and Semenkovich, 2012; Hadrich et al., 2018; Luo et al., 2019).

In summary, the present reveal the anti-diabetes effect of BefA in mammal for the first time, via increasing the number of islet β cells, reducing inflammatory reaction and apoptosis, improving liver lipid metabolism and restoring microbial diversity to normal level, which provides a new strategy for DM oral drug via inhibiting the progression of islet β cell destruction in diabetes (Fig. 7). However, a deeper understanding of the underlying mechanisms, and the use of engineered strain to replace the continuous administration of BefA is necessary to be further studied (Braat et al., 2006; Martín et al., 2014).

**Figure 7.**
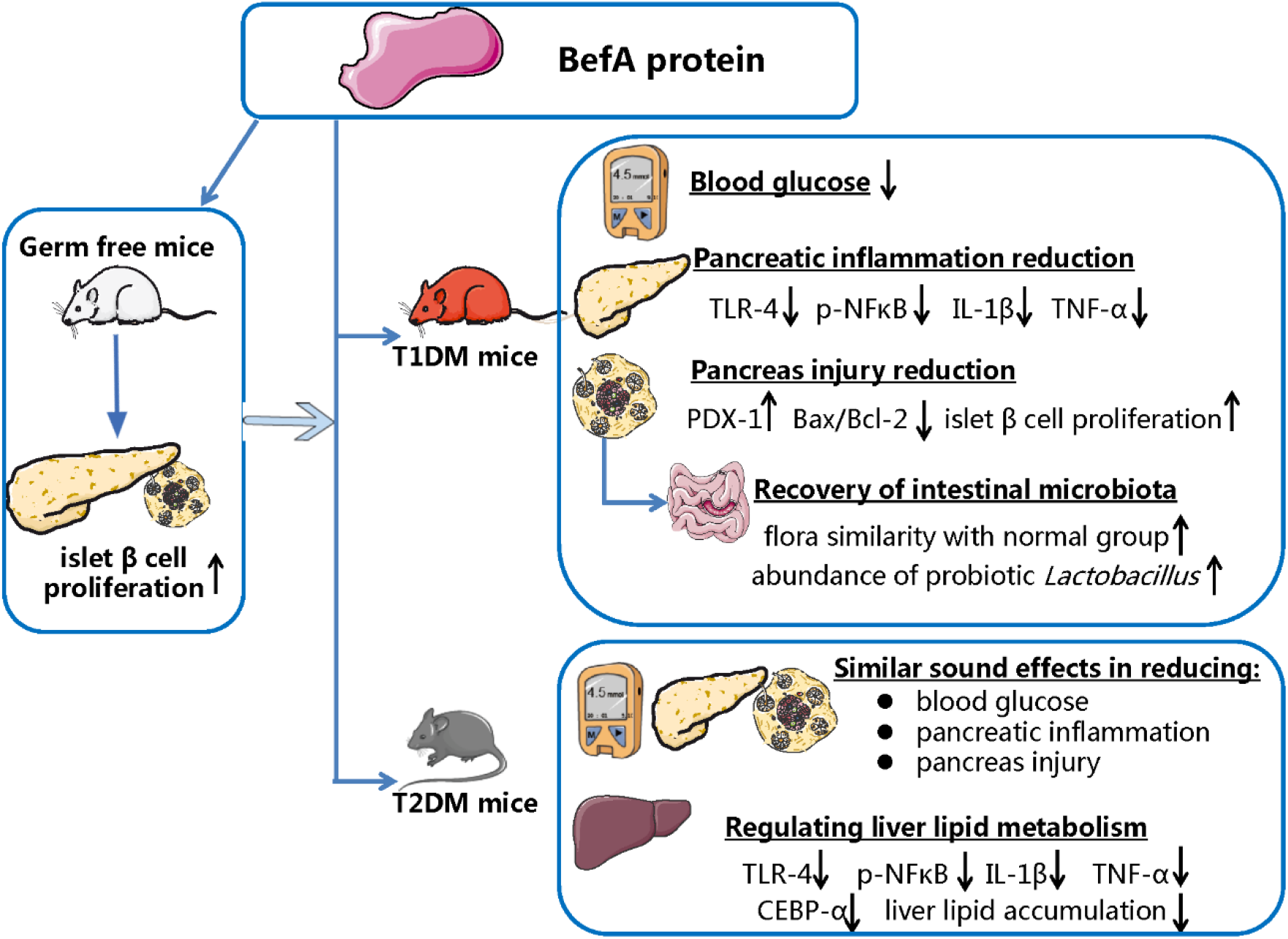
Mechanism diagram of BefA protein on diabetes.

## Materials and methods

### Construction of the BefA yield strain and protein purification

The BefA gene (M001_10165) was codon optimized for *Escherichia coli* BL21 to favour higher protein yield, and was synthesized with a histidine (His) tag (to facilitate the identification and purification of BefA protein), which was inserted into the prokaryotic expression vector pet 28C in Kingsy Biotechnology Co. (Nangjing, China) to form the recombinant plasmid pet 28C-BefA. Then, pet 28C-BefA was transformed into *Escherichia coli* BL21 strain to generate the BefA production strain of BL21-pet 28C-BefA.

To produce BefA protein, the BL21-pet 28C-BefA strain was cultivated in Luria-Bertani (LB) medium (Solarbio Life Sciences, China, L1010) with kanamycin ((50 μg/ml. Solarbio Life Sciences, China, K1030) at 37 °C. When the optical density value reached to 0.6-0.8, 1 mM of isopropyl β-D-thiogalactoside (IPTG, Solarbio Life Sciences, China, I8070) was added into culture medium to stimulate massive protein expression during the following 6-h cultivation. Then the culture medium was centrifuged at 8,000 g for 30 min to obtain the bacterial pellet, which was further used for ultrasonic disintegration to flow out bacterial proteins. BefA protein was purified with His-tag nickel beads (7Sea Biotech, China, PAN001-001C), and the purity and accuracy of BefA protein were detected by SDS-PAGE electrophoresis and Western-blotting, and the purified protein concentration was determined by BCA protein assay kit (Thermo Fisher Scientific, USA, 23227) according to the manufacturers guidelines.

### Construction and intervention of DM mice

To check whether BefA can affect the proliferation of Beta cells, newborn germ free mice (GF group; n=3), newborn SPF mice (GF group; n=3) and newborn germ free mice treated with 1 ng BefA/g body weight of BefA (GFB group; n=3) were used. For germ free mice, after the birth of mice born by female germ free mice in aseptic isolation, the sterilized were transferred into the aseptic isolation via isolation bin, and the BefA were given via oral gavage by skilled test staff.

For T1DM mice model, 8-week male wild-type C57BL/6 mice (purchased from SJA Laboratory Animal Co., Ltd, China) were housed in specific pathogen-free conditions with an optimum environment (12 h light/dark cycle with *ad libitum* access to standard laboratory chow and water, humidity 50 ± 15%, temperature 22 ± 2°C). After acclimating for 1 week, some measures are meant to avoid cage effects on the microbime, including separated mice individually in isolation bins, ensured the same diet and sterile padding, and changed gloves frequently when changing case. And injected intraperitoneally with streptozotocin (STZ, 50 mg/kg/d) for 5 consecutive days until the blood glucose concentration rose to 11.1 mM (199.8 mg/dl) or above and stabilized for 7 days, then mice were divided into 3 groups: (1) M group: T1DM model group treated with 0.9% physiological saline containing 0.01% gelatin administered intragastrically every other day for 14 times (n=15); (2) MB10 group: T1DM model group treated with 0.9% physiological saline containing 0.01% gelatin and 10 μg BefA administered intragastrically every other day for 14 times (n=15); (3) MB50 group: T1DM model group treated with 0.9% physiological saline containing 0.01% gelatin and 50 μg BefA administered intragastrically every other day for 14 times (n=15). Another 15 wild-type C57BL/6 mice were used as normal control group (C group). Within each group, all mice were used to test blood glucose level (once a week), body weight (once a week) and fecal microbiota structure (at week 7), in which 4 mice were sacrificed for pancreas Western-blotting analysis, 4 mice were sacrificed for pancreas qPCR analysis, the 3 mice were sacrificed for pancreas HE staining, immunehistochemical staining and immunofluorescent staining.

For T2DM mice model, 8-week-old male wild-type C57BL/6 mice were acclimated for 1 week, then fed with high fat diet (Research diets, USA, D12492) for 6 weeks combined with low-concentration of STZ (30 mg/kg) intraperitoneal injection until the blood glucose concentration rose to 11.1 mM (199.8 mg/dl) or above and stabilized for 7 days (Moreira et al., 2015; Helene, 2016; Ghiasi et al., 2015). Then mice were divided into 5 groups for 15 mice each: (1) C group: mice fed with laboratory chow diet as normal control group; (2) M group: mice treated with 0.9% physiological saline containing 0.01% gelatin administered intragastrically every other day for 14 times (n=15); (3) MB5 group: mice treated with 0.9% physiological saline containing 0.01% gelatin and 5 μg BefA (n=15); (4) MB20 group: mice treated with 0.9% physiological saline containing 0.01% gelatin and 20 μg BefA (n=15); (5) MB40 group: mice treated with 0.9% physiological saline containing 0.01% gelatin and 40 μg BefA (n=15). Within each group, all mice were used to test blood glucose level (once a week), body weight (once a week) and glucose tolerance (at week 10), in which 4 mice were sacrificed for pancreas and liver Western-blotting analysis, 4 mice were sacrificed for pancreas and liver qPCR analysis, 3 mice were sacrificed for pancreas immunehistochemical staining and liver oil red staining, 3 mice were sacrificed for pancreas immunofluorescent staining. The BefA concentrations used in T1DM mice model and T2DM mice model were determined by the pre-experiment based on the concentration previously used for zebrafish (Hill et al., 2016).

### Glucose tolerance test for DM mice

Mice were fasted for 12 h prior to the test. Glucose (1.5 mg/g) was injected intraperitoneally and blood glucose levels were measured at 0,30,60,90,120 min after the injection.

### Pathological histology

Pancreas sections were fixed in 4% paraformaldehyde, embedded in paraffin, cut into 5 μm sections and rehydrated by xylene and declining grades of ethanol for 5 min. Then, they were washed 3 times for hematoxylin and eosin (H&E) staining. Immunohistochemical and immunofluorescent tests were performed using anti-pancreas/duodenum homeobox protein-1 (PDX-1, Abcam, UK, ab47383) antibody and anti-insulin (Cell Signaling Technology, USA, #4590S) antibody. Frozen livers were sliced and Oil red O staining was performed as described earlier (Mariño et al., 2017; Ji et al., 2019) using Oil Red O Stain Kit (Lipid Stain)(SenBeiJia Biological Technology, China, BP-DL101). According to the instruction of the kit, the Oil Red O Solution was added dropwise onto the tissue for a 5-10 min incubation period. Excess staining buffer was removed with 85% Propylene Glycol. The tissues were washed with distilled water and counterstained with hematoxylin.

### Western blotting analysis

Tissues were lysed in radio immunoprecipitation assay (RIPA) lysis buffer (Solarbio Life Sciences, China, R0010) and centrifuged at 8,000 g for 15 min at 4 °C after sonication on ice. The supernatants were collected and the protein concentrations were measured by BCA assay. Equal amounts of sample (60 mg/lane) were heat-denatured in loading buffer and separated via 10% gel electrophoresis (SDS-PAGE) followed by transferred onto PVDF membrane (Millipore, Germany, IPVH00010). Then the membrane was blocked with 5% skim milk-TBST solution (20 mM Tris-HCl (pH 7.6), 127 mM NaCl, 0.1% Tween 20) for 1 h at room temperature (Chen et al., 2019). After washed 3 times by TBST, the samples were incubated overnight with primary antibodies directed against anti-β-actin (ABclonal, USA, AC026), anti-His tag (Solarbio Life Sciences, China, K200060M), anti-toll-like receptor-4 (TLR-4) (Santa Cruz Biotechnology, USA, sc-293072), anti-nuclear factor kappa-B (NFκB) (Abcam, UK, ab32360), anti-phosphorylated nuclear factor kappa-B (p-NFκB) (Santa Cruz Biotechnology, USA, sc-101751), anti-Bcl-2-associated X protein (Bax) (Cell Signaling Technology, USA, # 5023S), anti-B cell lymphoma-2 (Bcl-2) (Cell Signaling Technology, USA, #3498S) and anti-CCAAT enhancer binding protein-α (CEBP-α) (Cell Signaling Technology, USA, #2295S) at 4 °C. The membranes were then washed 3 times by TBST and incubated with horseradish peroxidase (HRP)-linked anti-rabbit IgG or anti-mouse IgG antibodies at room temperature for 1 h. Membrane-bound immune complexes were detected by enhanced chemiluminescence system (Thermo Scientific, USA). Quantification was performed by densitometric analysis using the Image J software (NIH). All Western blotting experiments of each protein were carried out with four experimental replications, see in supplementary Fig. S1.

### RNA extraction and q-PCR

Mouse pancreas and livers were homogenized in Trizol reagent (Life Technologies, USA, 15596026) prior to RNA extraction, PCR primers were designed using Primer 5.0. q-PCR amplification was performed by ABI 7900HT fast real-time PCR system (Applied Biosystems, USA). The reaction mixture contained 10 μL of SYBR^®^ Primer EX Taq II (Takara, Japan, RR420A), 0.4 μL ROX Reference Dye (50×) (Takara, Japan, RR420A), 1.0 μL DNA template and 0.8 μL each of the primers (final concentration was 0.4 μM), with 7 μL Milli-Q H_2_O. The q-PCR condition was: start at 95 °C for 10 min, followed by 40 cycles of degeneration at 95 °C for 30 s, annealing at 60 °C for 30 s and extension at 72 °C for 30 s. Relative levels (fold change) of the target genes were normalized against a housekeeping gene (GAPDH) and analyzed by the 2^−(△△Ct)^ method (For specific primers, see Table 1).

**Table 1.**
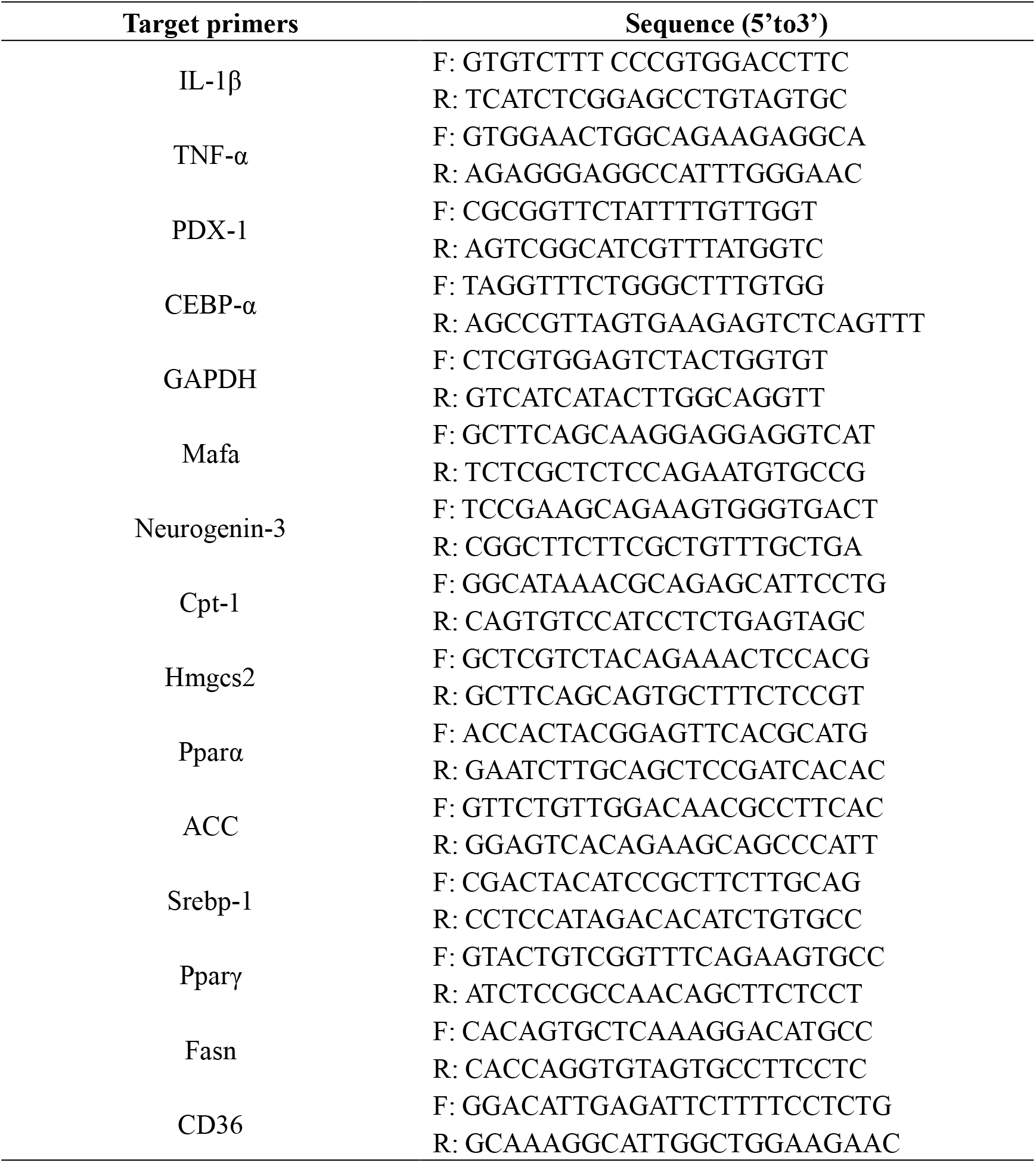
Primers for amplifying different 469 genes via q-PCR.

### High-throughput sequencing analyses

Feces of mice were collected and were stored at -80 °C. Bacterial genomic DNA from feces was obtained using DNA extraction kit (Tiangen, China, DP302). DNA samples were amplified targeting the V3-V4 region of bacterial 16S rRNA gene using 338F/806R primers (Geller et al., 2017). Bioinformatics analysis was performed with UPARSE software version 7.0.100 (http://drive5.com/uparse/) using the UPARSE-operational taxonomic units (OTU). Sequences with ≥97% similarity were assigned to the same OTUs. Weighted unifrac distance analysis was performed using the quantitative insights into microbial ecology (QIIME) software package version 1.9.1 (http://qiime.org/; QIIME Development Team) and LEfSe (Linear discriminant analysis effect size) method was used to analyze the bacteria with significant differences among C, M, MB10 and MB50 groups (GenBank accession no. PRJNA637680) (Chen et al., 2018).

### Statistical analysis

Data handling, analyses and graphical representations were performed using Graphpad Prism version 7.0 (GraphPad Software, Inc. USA). Values are shown as mean ± SD. Statistical significance was determined using one-way or two-way ANOVA and were annotated using the international convention related to the statistical representation.

## Conflicts of interest

The authors declare no conflict of interest.

## Data availability

The authors confirm that the data supporting the findings of this study are available within the article and its supplementary materials. Raw sequences have been deposited in the GenBank under accession number PRJNA637680.

## Ethics declarations

All experimental procedures involving mice were approved out according to protocols by Laboratory Animal Ethics Committee of Nanchang Royo Biotech Co., Ltd (RYL2017042901).

## Authors’ contributions

TC and JL conceived the study and participated in its design and coordination. HW, JW, HH involved in sample collection and performed the experiments and bioinformatics analyses. TC, HW and JL revised the manuscript and all authors approved the final manuscript.

## Acknowledgements

We are grateful to the scientific research staff of Institute of Translational Medicine of Nanchang University for their contributions to this study.

## Funding Information

This work was supported by grants from the National Natural Science Foundation of China (31560264 to TC), the Excellent Youth Foundation of the Jiangxi Scientific Committee (20171BCB23028 to TC), the Science and Technology Plan of the Jiangxi Health Planning Committee (20175526 to TC), the Science and Technology Project of Jiangxi (20181BBG70028 and 20181BCB24003 to TC).

---

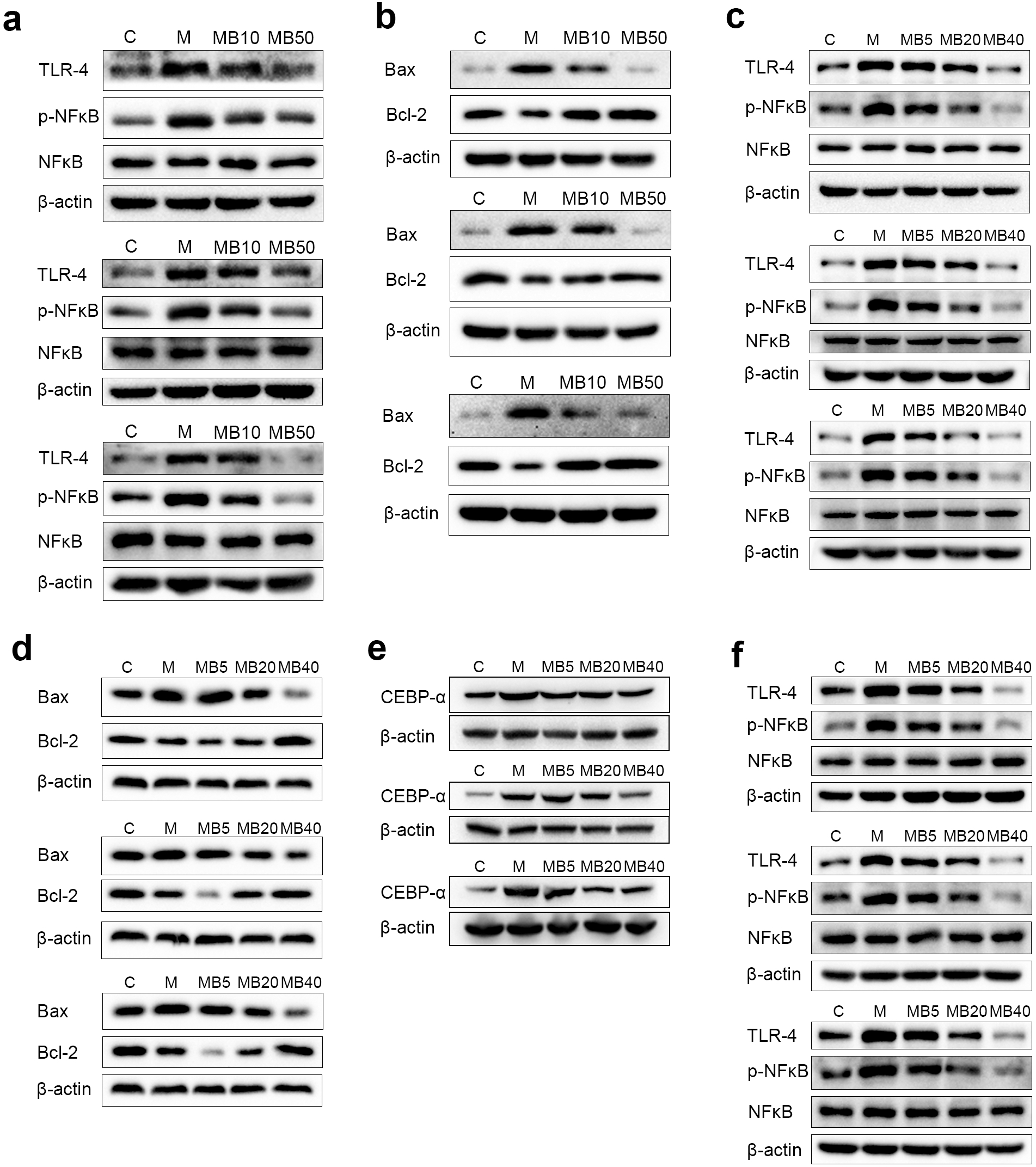

## Notes

### Competing Interest Statement

The authors have declared no competing interest.

